# Mutating the interprotofilament interface allows microtubules to assemble in GDP

**DOI:** 10.64898/2026.07.09.737542

**Authors:** Yean Ming Chew, Robert A. Cross

## Abstract

Microtubule dynamic instability, driven by GTP turnover, allows microtubules in cells to reorganise themselves adaptively. GTP promotes the assembly of tubulin into microtubules, but exactly how it does so is controversial. In some models, GTP mainly supports the assembly of tubulin into single protofilaments, as in prokaryotic tubulins. In others, GTP mainly supports the formation of lateral bonds between protofilaments. To investigate, we mutated the interprotofilament interface in human α1bβ3 and α1bβ4b tubulins, whose sequences diverge markedly in this region. We find that transplanting the α1bβ3 M-loop or its binding pocket into α1bβ4b tubulin creates tubulins that assemble in 1 mM GDP. We accordingly propose that GTP- and GDP-tubulins are captured equivalently at the tips of microtubules, but then differentially retained, based on their differing abilities to form stable interprotofilament bonds. This biased retention mechanism allows mosaic lattices to be built and dynamic instability to be tuned.

**One sentence summary:** Microtubules grow not by biased capture of GTP tubulins, but by biased retention of tubulins that form stable interprotofilament bonds.

## INTRODUCTION

Microtubules (MTs) are polymeric, energy-dissipating GTPase molecular machines that are essential for eukaryotic life. Polymerisation of GTP-tubulin activates its GTPase, so that in growing MTs a cap of newly-recruited GTP-tubulin comes to overlie an unstable GDP-core (Mitchison and Kirschner, 1984; Roostalu et al., 2020). Breaching the GTP-tubulin cap exposes the unstable GDP-tubulin core lattice, triggering catastrophic disassembly (Brouhard and Bement, 2015). The resulting cycle of dynamic instability, driven by GTP turnover, allows MTs in cells to act as adaptive rails, actuators and sensors (Mitchison, 2026).

Until recently, models of dynamic instability envisaged that GTP drives tubulin molecules in solution into a straight conformation that is more favourable for polymerisation than the curved conformation of GDP-tubulin, and that this causes recruitment of GTP-tubulin into the MT lattice via a conformational selection mechanism (**Fig. 1A**). However, there is now firm evidence that GTP-tubulin and GDP-tubulin have similar conformations in solution (Wagstaff et al., 2023; Zhou et al., 2023), pointing away from a conformational selection mechanism and towards an induced fit mechanism, in which incoming subunits are switched into a polymerisation-friendly conformation by the polymerisation reaction (**Fig. 1B**). Polymerisation-coupled conformational switching at the level of single PFs is well-evidenced for prokaryotic tubulins, which assemble only into single PFs, and might also operate for PFs at the splayed tips of eukaryotic microtubules (Erickson, 2019). In eukaryotic MTs, it is further possible that structural switching might be actuated partially or entirely by lateral bonding between nearest-neighbour αβ tubulin heterodimers in the growing lattice (Rice et al., 2008).

**Figure 1.**
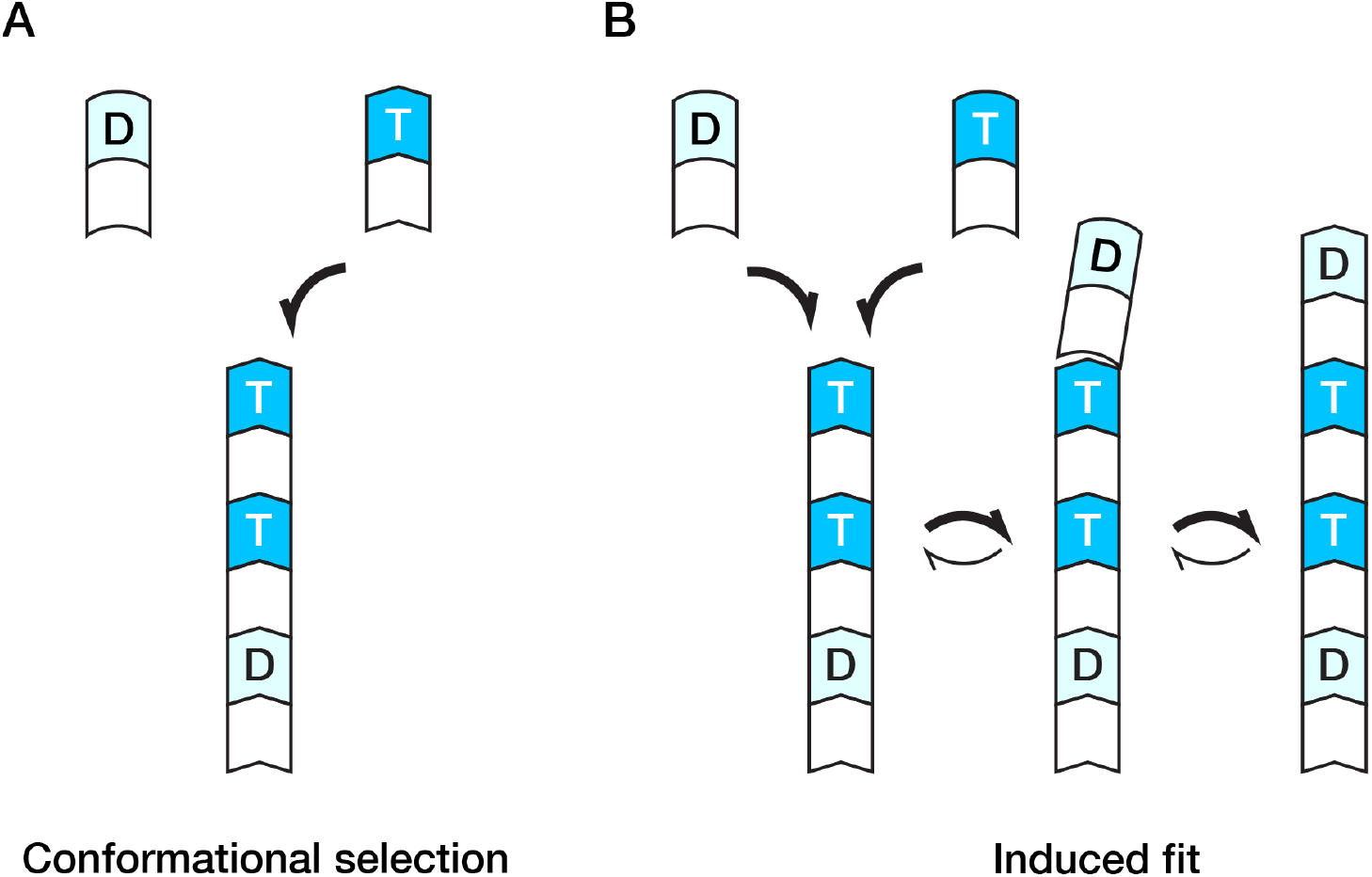
Polymerisation-coupled structural switching at the level of a single protofilament. **(A)** In a conformational selection mechanism, unpolymerized tubulin subunits can have 2 conformations, one stabilised by GTP (T) and the other by GDP (D). The tip of the growing polymer selects the polymerisation-friendly GTP conformation. **(B)** In an induced fit mechanism, unpolymerized GTP- and GDP-tubulin subunits are both in the polymerisation-unfriendly conformation. Incorporation into the polymer switches them into the polymerisation-friendly conformation, so that the PF tip catalyses its own growth. GTP may facilitate, but is not required for, this structural switch. For simplicity the polymerisation-coupled conformational switch is shown operated fully by the initial capture reaction. In fact full switching may occur only upon addition of a further, overlying subunit (Erickson, 2019).

Whilst GTP is indispensable for MT dynamic instability, it is not clear that it is required for MT growth. There is evidence that GTP-tubulin and GDP-tubulin can co-polymerise (Valiron et al., 2010), that GTP and GDP can exchange at the tips of MTs (Piedra et al., 2016), and that pure GDP-tubulin at very high concentrations can polymerise on to the minus ends of stabilised MT seeds (Bagdadi et al., 2024). Further, recent work suggests that the main action of GTP may be to make tubulin more flexible (Shred et al., 2025), so as to facilitate interprotofilament (interPF) bonding. This prompted us to ask, can mutation of the interPF interface promote the assembly of GDP-tubulin into MTs?

## RESULTS AND DISCUSSION

To investigate, we expressed human α1bβ3 and α1bβ4b tubulins in insect cells, tagged with an 8-his on the α tubulin C-terminus and a FLAG on the β tubulin C-terminus (Chew and Cross, 2023; Minoura et al., 2013). Eukaryotic tubulins form interPF contacts by engaging the α and β M-loops of each tubulin heterodimer with a recognition surface consisting of the H1’-S2 and H2-S3 loops on the lateral nearest-neighbour in the lattice (Prota et al., 2023) (**Fig. 2**). We focus here on the β tubulin M-loop interface. Excepting the C-terminal tubulin code sequences, surface-exposed sequence differences between human β-tubulin isotypes cluster at this interface (**Fig. 2A**)(Wood and Moore, 2025).

**Figure 2.**
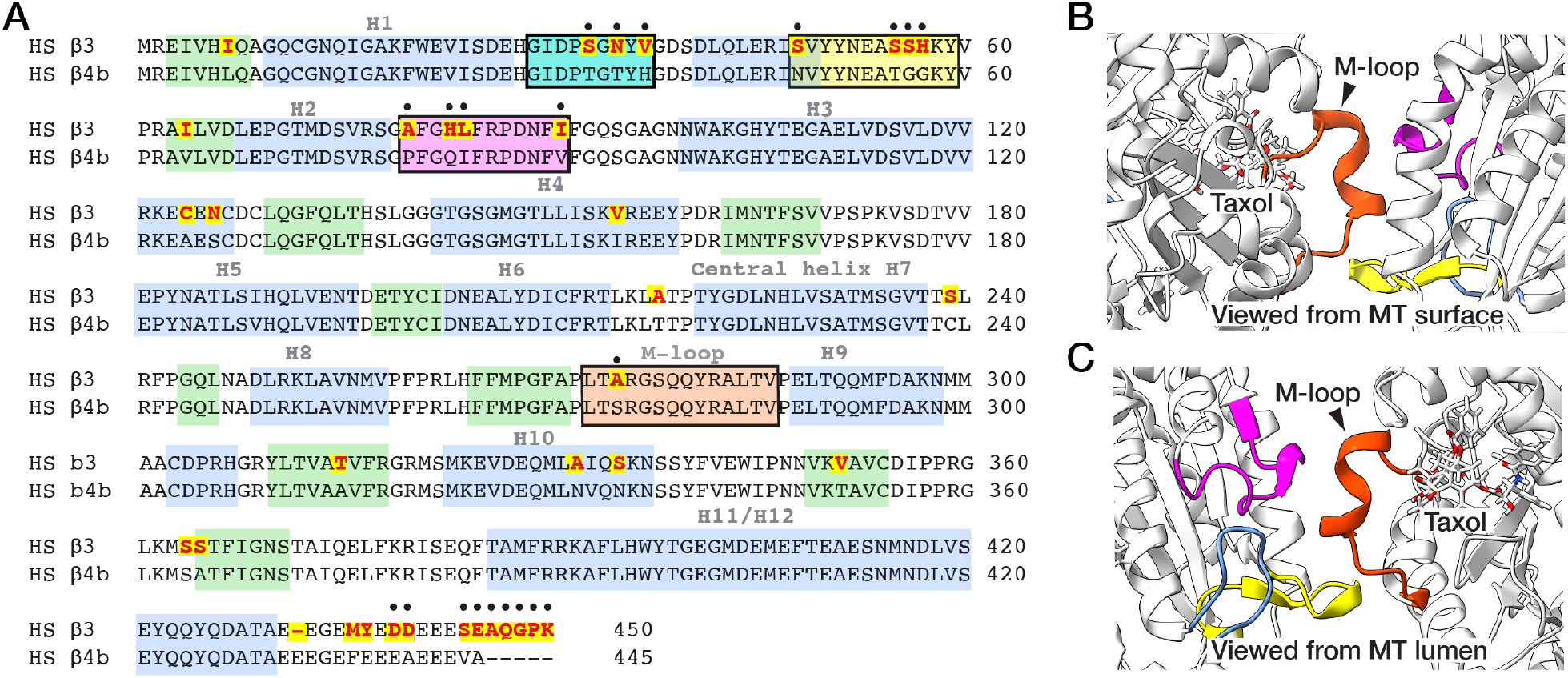
Beta tubulin M-loop interface. **(A)** Human β3 and β4b tubulin sequences are very similar, except at their C-termini and in their β M-loop interface. Divergent residues are highlighted in red on yellow. Divergent residues that are surface exposed (black dots) cluster in the M-loop binding pocket. **(B)** β-tubulin M-loop interface in 6WVM.pdb (Debs et al., 2020). The M-loop (orange) engages a surface composed of H1’-S2 (yellow) and H2-S3 (magenta) loops. **(C)** Same interface viewed from the MT lumen. The taxol site abuts the M-loop.

In human α1bβ3 to α1bβ4b microtubules, clusters of divergent residues in the lateral interacting sites are associated with altered dynamics and altered sensitivity to taxol (Chew and Cross, 2023). Taxol binds close to the lateral interaction interface (**Fig. 2B, C**). Taxol is not only an important chemical biological tool but also a critically-important therapeutic. Taxol stabilises human microtubules but is less effective on β3 tubulin. Mutating the lateral interaction surfaces can potentially illuminate both the mechanisms by which tubulin assembles, and the role of lateral interactions in determining the binding and microtubule-stabilizing action of taxol. Accordingly we engineered two sets of mutants, one set that swapped the M-loop binding pocket in β tubulin between α1bβ3 to α1bβ4b isotypes and vice versa, and another set that swapped the M-loop itself, by making point mutations at residues 218 and 275.

### Mutation of the M-loop interface profoundly affects MT assembly

As MTs grow, M-loop interactions are thought repeatedly to form and break (Gudimchuk and McIntosh, 2021). Even after the M-loops engage stably, they retain some flexibility, allowing the tube to articulate (Amos, 2010), whilst tending nonetheless to adopt preferred conformations that constrain the angles between PFs and the PF number (Brouhard and Rice, 2018). Under our conditions, wild-type single isotype α1bβ4b MTs and α1bβ3 MTs in 1 mM GTP show dynamic instability at both ends of their GMPCPP seeds (**Fig. 3B**). α1bβ3 MTs depolymerise about 2x faster than α1bβ4b MTs (Chew and Cross, 2023). Counterintuitively, we find that swapping the M-loops or their binding pockets between α1bβ3 and α1bβ4b tubulins *amplifies* this difference in lattice stability. Lattices built from α1bβ3 tubulin with a β4b M-loop or binding pocket are less stable than those built from WT α1bβ3. Lattices built from α1bβ4b tubulin with a β3 M-loop or binding pocket are more stable than those built from WT α1bβ4b. (**Fig. 3B**).

**Figure 3.**
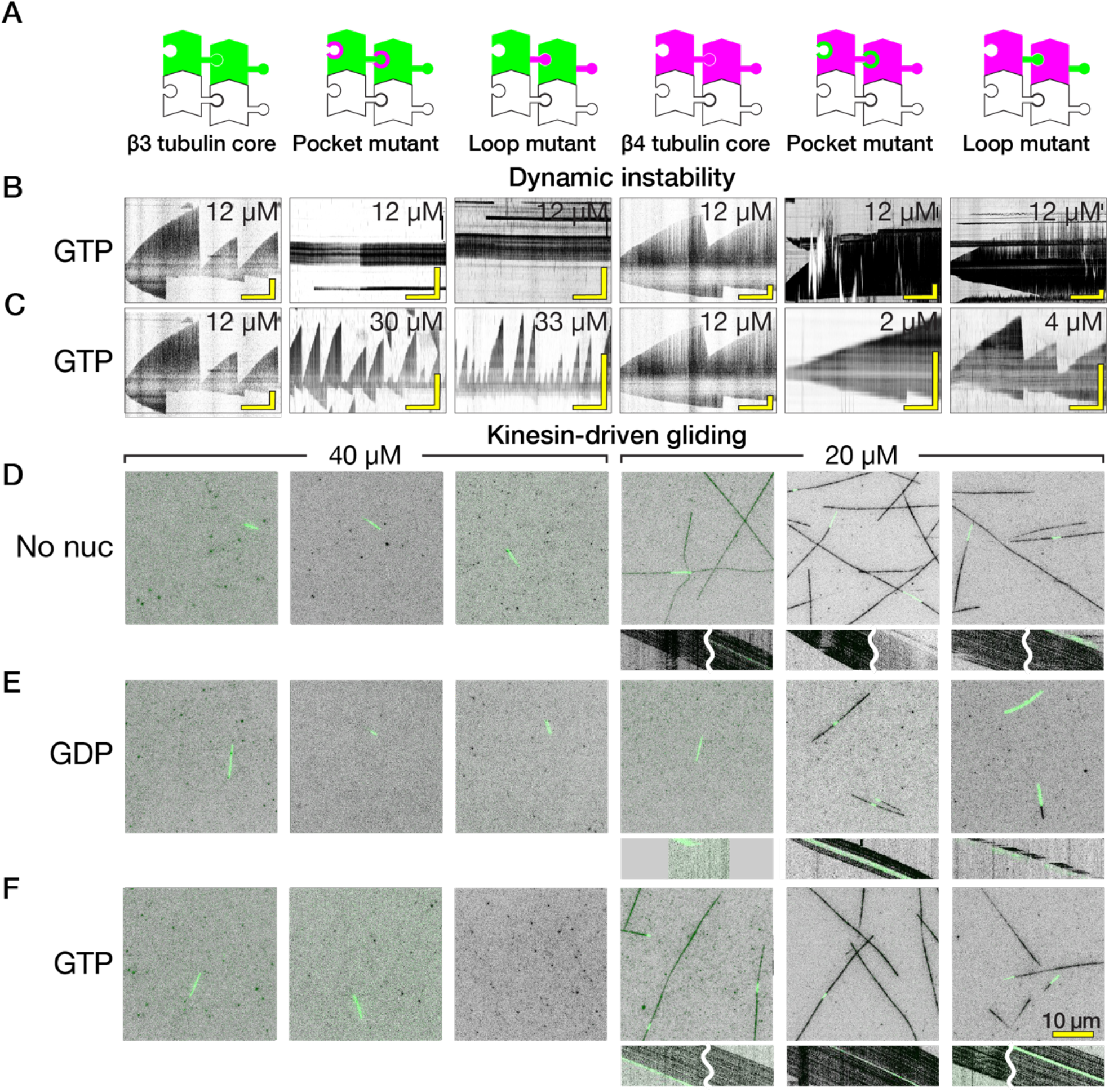
Beta M-loop interface mutants radically influence MT assembly. **(A)** Cartoons of WT and chimeric tubulins, viewed from surface, MT plus end up. **(B)** Kymographs comparing the dynamic instability of single isotype and mutant single isotype MTs at 12 µM tubulin in 1 mM GTP over 30 minutes of observation. WT α1bβ3 and α1bβ4b MTs show dynamic instability, whilst α1bβ3 with an α1bβ4b pocket or loop does not assemble onto the seeds and α1bβ4b with an α1bβ3 pocket or loop shows unbounded growth. Vertical scale bar is 5 µm in every case. (**C**) By adjusting the tubulin concentrations as shown, dynamic instability is observable for both WT and mutants. Darkfield imaging of unlabelled tubulins. **(D-F)** MT gliding on KIF5B surfaces. Green: GMPCPP stabilised pig brain seeds; black: MT extension. Kymographs beneath each frame illustrate the rates of kinesin-driven gliding. Some kymographs are split (wavey white line) to display both MT ends (**D**) Assembly with no added nucleotide. α1bβ3 constructs are unstable under these conditions. α1bβ4b WT and mutant MTs are robust and trackable in gliding assays. **(E)** Assembly with added 1 mM GDP. α1bβ4b chimerae assemble into MTs that glide at rates similar to WT. **(F)** GTP-assembled MTs glide similarly.

### InterPF interface mutants can assemble in 1 mM GDP

Our finding that chimeras with an α1bβ4b core and an α1bβ3 M-loop or binding pocket form hyperstable lattices suggests that adjusting the β tubulin interPF interface by mutagenesis can overcome a barrier to polymerisation set by the ability of each subunit to connect to its lateral nearest-neighbours (Cleary and Hancock, 2021; Gudimchuk and McIntosh, 2021; Kalutskii et al., 2025) We therefore asked, if lateral bonding is made sufficiently favourable, can the GTP requirement for MT growth be overcome entirely, allowing MTs to be built from GDP-tubulin? For these experiments, we triggered assembly not by the usual method of adding GTP, but by adding GMPCPP seeds to the tubulin solution and raising the temperature to 37° C. Tubulins were buffer-exchanged into GTP-free KPEM buffer by gel filtration as the last step in their preparation (see *Methods*), so that the assembly mix in these experiments contained a maximum of one exchangeable GTP molecule per β tubulin. Under these conditions, without added nucleotide, WT α1bβ4b tubulin and our α1bβ4b-based chimeras do assemble into microtubules, whereas WT α1bβ3 and its chimeras do not. This suggests that for α1bβ4b tubulin, but not α1bβ3 tubulin, micromolar concentrations of co-purifying GTP are sufficient to drive assembly. To investigate further, we added 1 mM GDP. Adding 1 mM GDP blocked assembly of the WT α1bβ4b, but remarkably, supported assembly of both α1bβ4b with a β3 M-loop binding pocket, and of α1bβ4b with a β3 M-loop, albeit to a lesser extent (**Fig. 3B**). The GTP- and GDP-assembled MTs (stabilised by glycerol) glide at similar rates on kinesin surfaces (**Fig. 3C**, kymographs).

### Taxol-induced lattice expansion depends on the interPF interface

Adding taxol assembles all the tubulins examined here (**Fig. 4**). Taxol binds ∼4 orders weaker to unpolymerized tubulin than to polymerised tubulin (Prota et al., 2023), consistent with its main action being to strengthen interPF connectivity (Manka and Moores, 2018). Taxol can polymerise brain GDP-tubulin (Diaz and Andreu, 1993) and can modulate tubulin conformation, including by expanding the axial spacing of tubulin dimers within the GDP-lattice (Kamimura et al., 2016; Lucena-Agell et al., 2026), isotype-specifically (Chew and Cross, 2023). Taxol-induced lattice expansion accelerates MT gliding on a surface of *Drosophila* kinesin-1 (Chew and Cross, 2023). MTs gliding on a human KIF5B surface respond similarly (**Fig. 4**). As we reported earlier, taxol accelerates the gliding of WT α1bβ4b MTs, but not WT α1bβ3 MTs, indicating α1bβ4b MTs are expanded by taxol, whereas α1bβ3 MTs are not. Swapping the β4b M-loop into β3 tubulin, by mutating only two residues at position 218^th^ and 275^th^, allows taxol to accelerate MT gliding (**Fig. 4**), suggesting taxol binding is potentiated. Conversely, swapping the β4b M-loop binding pocket for its β3 equivalent slows gliding (**Fig. 4**), suggesting taxol binding is reduced (Bozdaganyan et al., 2025).

**Figure 4.**
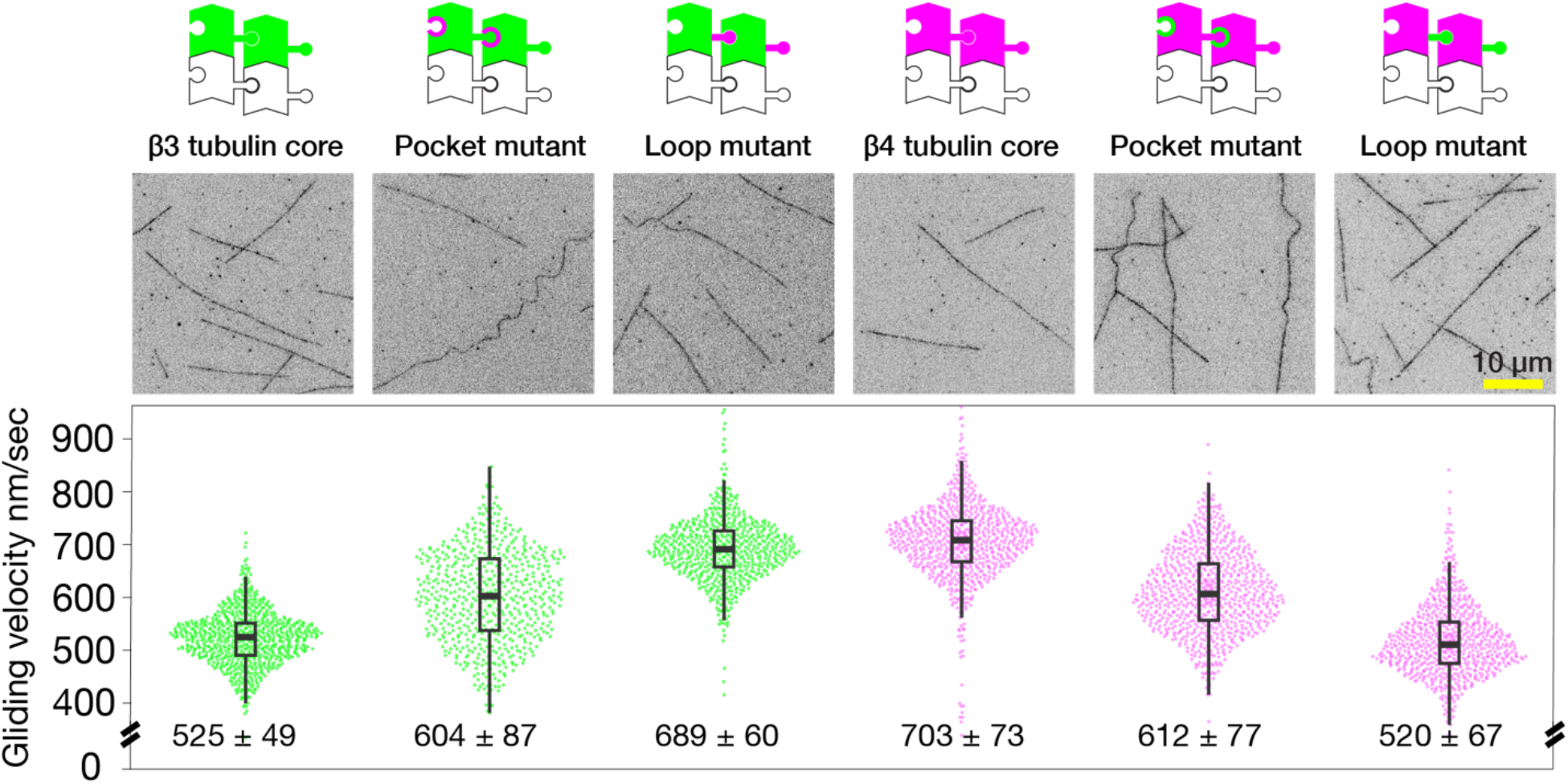
Effects of taxol on kinesin-driven MT gliding depend on the interPF interface. All tubulins assemble in 10 µM taxol. Gliding velocities of single isotype microtubules in 2 mM ATP and 10 µM taxol, on a surface of human full-length KIF5B (mean ± SD, n = 782, 616, 665, 746, 781 and 712, n is instantaneous velocity). WT α1bβ3 MTs are not expanded by taxol and glide more slowly than WT α1bβ4b MTs. MTs built from chimeric tubulins glide at intermediate rates, suggesting taxol binding to β3-based chimeras is increased, and that to β4b-based chimeras is decreased. In contrast, mutations in M-loops fully reverse taxol responses.

### Hyper-assembler mutants can recruit hypo-assembler mutants into a mosaic lattice

To further probe the role of interprotofilament annealing in MT polymerisation, we asked whether hyper-assembler tubulins can drive hypo-assembler tubulins to polymerise. We tracked incorporation of each isotype into the lattice using tris-NTA fluorophores, which bind tightly to the his-tags. Indeed, we see that tubulins that do not otherwise assemble are recruited into the lattice by our hyper-assembler mutants (**Fig. 5A, B**). The resulting mosaic lattice depolymerises markedly more quickly following glycerol washout than that built purely from the hyper-assembler (**Fig. 5A, B** kymographs).

**Figure 5.**
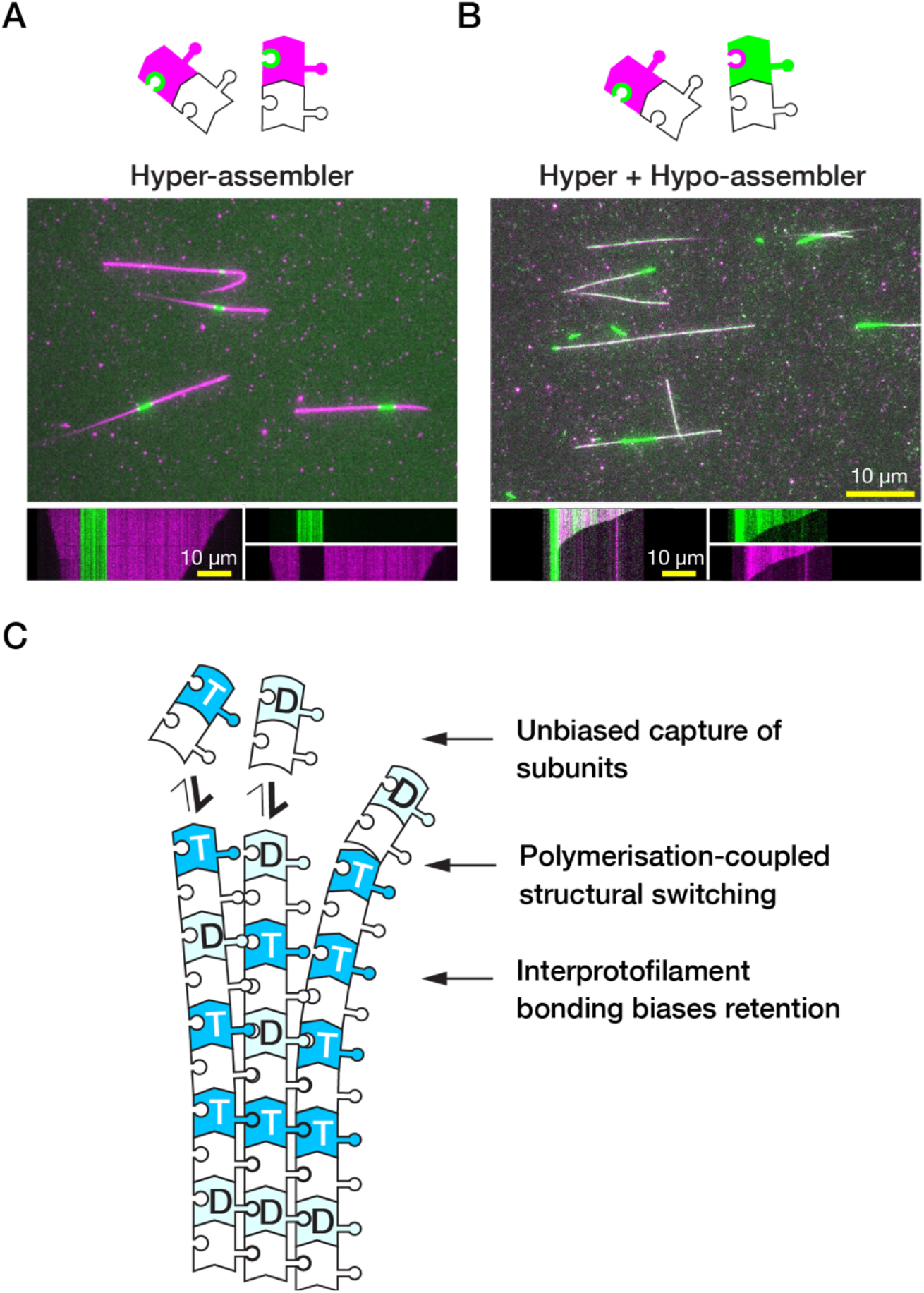
Hyper-assembler tubulin can recruit hypo-assembler tubulin into a mosaic lattice. **(A)** β4b-tubulin with β3 M-loop binding pocket (magenta) assembles readily onto GMPCPP brain tubulin seeds (covalently labelled in green). **(B)** A 1:1 mix of hyper-assembler (magenta) and hypo-assembler (green) mutant tubulins recruits both tubulins into a mosaic lattice. GMPCPP seeds again covalently labelled in green. **(C)** Proposed biased retention model. Polymerisation-coupled structural switching occurs at the PF level for both GDP-and GTP-tubulin but retention by the core lattice is biased towards GTP-tubulin because it more readily forms stable interPF bonds.

Our data show that mutating the inter-PF interface can erode the functional contrast between GTP- and GDP-tubulin, allowing GDP-tubulin to form a stable lattice. Mutations at other positions, including in the active site (Estévez-Gallego et al., 2025) and flanking the central H7 helix (Ye et al., 2020) have been shown also to favour MT assembly, but are not reported to overcome the requirement for GTP (Mondal et al., 2025). By swapping the M-loop or its recognition interface from β3 into β4b tubulin, we have overcome a kinetic barrier to tubulation that is clearly an evolved feature of both natural isotypes. This points strongly to a model (**Fig. 5C**) in which MTs provisionally recruit both GTP and GDP-tubulin, but then preferentially retain GTP-tubulin, based on its greater ability to make stable lateral connections. Most current models posit a zone at the tips of growing MTs in which the longevity, curvature and flexibility of exposed PFs (McIntosh, 2025) determine the probability of their tubulin molecules ultimately incorporating into the lattice. Recent work indicates that the GTP-dependent bundling of exposed PFs (Kalutskii et al., 2025) is a critical step along this pathway. Our data do not speak to the detailed structure of MT tips, but they do imply a zone into which tubulin molecules are provisionally recruited and then retained or rejected depending on their ability to connect stably to their lateral nearest-neighbours.

Our finding that tubulin retention by the lattice can be increased by mutagenesis to the point that GTP becomes unnecessary does not imply a diminished role for GTP in the assembly of WT tubulins, or for any of the many other allosteric interactors that regulate MT dynamics. Rather, it clarifies the reactions on which these effectors act and emphasises that evolution has tuned the actions of GTP and GDP on the interPF interface to achieve regulable dynamic instability. Allosteric interactors other than GTP, including different tubulins, post-translational modifiers, MAPs, polymerases, depolymerases, motors, and drugs can be pivotal in regulating the dynamic instability (Chew and Cross, 2025; Cross, 2019) and lattice structure (Guyomar et al., 2022; Romeiro Motta et al., 2023) of natural MTs. But among these effectors, GTP is the only one cleaved by the MT. GTP fuels dynamic instability and is indispensable for MT assembly from WT tubulin at physiological concentrations (Ayukawa et al., 2021; Mondal et al., 2025). Our finding that the requirement for GTP can be overcome by mutagenesis of the interPF interface directly demonstrates that tubulin itself is a central allosteric effector for tubulin polymerisation. It seems possible that all allosteric effectors of tubulin assembly, including GTP, actuate a common assembly switch (Buey et al., 2006; Wagstaff et al., 2023) whose operation in modern tubulins is potentiated by interPF connectivity.

## Materials and Methods

### Buffers and reagents

Buffer BRB 80 was composed of 80 mM PIPES, 1 mM EGTA and 1 mM MgCl_2_, pH adjusted to 6.8 with KOH pellets. Buffer KPEM was made up of 100 mM PIPES, 2 mM EGTA and 1 mM MgSO_4_, pH adjusted to 6.9 with KOH pellets. Our oxygen scavenger of choice was glucose oxidase and catalase (GOC). The components were stored separately at 100× concentration at -20°C and constituted to 1× immediately before the experiment. The final concentrations used in assays were 4.5 mg/mL glucose, 0.2 mg/mL glucose oxidase, 35 μg/mL catalase.

### Tubulin and kinesin constructs

An L21 leader sequence (AACTCCTAAAAAACCGCCACC) was placed 5’ of tandem α and β tubulin sequences to enhance protein expression α and β tubulins were C-terminally tagged with 8x his (HHHHHHHH) and FLAG (DYKDDDDK) sequences, respectively, with a linker GGSGG in between, as previously described (Chew and Cross, 2023; Minoura et al., 2013). Expression of both α and β tubulins was driven by polyhedrin promoters.

Assembly of both α and β tubulin genes used the biGBac system (Weissmann et al., 2016). Individual α and β tubulin genes obtained from GeneArt were amplified using pLIB_for and pLIB_rev primers. These amplicons were subsequently cloned into pLIB vector which contains a polyhedrin promotor and a SV40 terminator using Gibson Assembly. The gene expression cassettes which consist of a promoter, α or β tubulin gene, and a terminator were amplified again using Cas_I_for and Cas_I_rev primers for α tubulin gene; and Cas_II_for and Cas_ω_rev primers for β tubulin gene. These α and β tubulin gene expression cassettes were then joined together via Gibson Assembly into a pBIG plasmid. These pBIG-α-β tubulin constructs were propagated in *Escherichia coli* (*E. coli*) DH5α cells. Positive clones were isolated using antibiotics and integrated into baculovirus genome via the Tn7 transposition sites in *E. coli* DH10EmBacY. Colonies for bacmid generation were selected using blue-white screening.

Bacmid DNA was isolated from an overnight *E. coli* culture using MidiPrep reagents (Qiagen) and overnight DNA precipitation at -20°C instead of using a spin column. Bacmids were pelleted for 15 minutes at 14, 000 × g, rinsed in 70% ethanol and pelleted again. The pellets were air-dried and resuspended in water. PCR verification was done using M13 forward and reverse primers to check for successful integration of the genes. Bacmids were stored at 4°C and used within two weeks.

Human KIF5B (gene KIF5B; NP_004512.1) was C-terminally tagged with twin-Strep tag sequence (SAWSHPQFEK<u>GGGSGGGSGG</u>SAWSHPQFEK, linker underlined) in pFastBac backbone (GeneArt). Expression was driven by a polyhedrin promoter. The Bacmid was obtained in the same way as for tubulin.

### Baculovirus production and protein expression

Sf9 insect cells were maintained at 28°C with shaking at 120 rpm in Ex-Cell 420 (Merck, 14420C) medium, supplemented with 100 U/mL penicillin-streptomycin, split to a density of 0.5 × 10^6^ cell/mL when reaching 2 × 10^6^ cell/mL.

Transfection was achieved by seeding 2 mL culture at 0.5 × 10^6^ cell/mL onto a 35 mm petri dish, incubated for 15 minutes at room temperature to allow adherence. In the meantime, a transfection reaction was prepared by mixing 2.5 μg of bacmid into 200 μL of medium, followed by 6 μL of FuGene HD. This mix was incubated for 15 minutes and then dripped over the seeded cells, which were subsequently incubated for about 3 to 4 days at 28°C without shaking. P1 virus was harvested by spinning down at 750 × g for 5 minutes to remove cell debris.

P2 virus was generated by adding 1 volume of P1 virus to 100 volumes of cell culture with a density of 1.0 × 10^6^ cell/mL (typically 0.5 mL of P1 virus to 50 mL of culture. The culture was clarified by centrifugation at 750 × g for 5 minutes. The supernatant was filtered through a 0.45 μm PVDF syringe filter and kept at 4°C for storage. P3 virus was used for protein expression and was produced similarly.

Protein expression was done by adding 30 mL of P3 virus into 1 L of culture when it reached a density of approximately 2 × 10^6^ cell/mL. For tubulin purification, 4 L of culture was normally used to allow enough yield for biochemical assays. Only 1 L of culture was needed for kinesins. The cells were harvested usually 52 hours post-infection by spinning down at 750 × g for 20 minutes at 4°C. The pellets were rinsed with ice-cold PBS and spun down again. The pellets were flash-frozen in liquid nitrogen and kept at -80°C for storage.

### Tubulin purification

All purification steps were performed at 4°C or on ice, unless otherwise stated.

Typically, 4 L of culture was used for every purification to produce enough protein for biochemical assays. An equal volume of lysis buffer composed of KPEM buffer supplemented with 0.5 M 3-(1-pyridinio)-1-propane sulfonate, ∼250 unit/mL Benzonase® nuclease, 1 mM DTT, 1 mM PMSF, 0.05% CHAPS, 25 mM imidazole, 1% glycerol, 1 mM ATP and 1 mM GTP was added to the cell pellets and the mix incubated on ice for about 15 minutes. The mixture was homogenised with a douncer with about 50-60 strokes. The lysed cells were clarified by spinning down at 200, 000 × g for 35 minutes (Thermo Scientific, T-865 fixed-angled rotor). The supernatant was loaded onto a 5 mL HisTrap column using an ÄKTA Purifier.

Pooled fractions after Ni-NTA purification were diluted 2× in KPEM to lower KCl concentration to 125 mM to improve tubulin binding. Subsequently, 5 mL of FLAG resin (A2220, Milllipore) was added and the mixture rolled for an hour. The slurry was then packed into a column. The resin was washed with 5 × volume of KPEM supplemented with 125 mM KCl, 1 mM ATP and 1 mM GTP. Tubulin was eluted with KPEM supplemented with 250 μg/mL FLAG peptide (F3290, Millipore) and 125 mM KCl, 1 mM ATP and 1 mM GTP. The eluates were pooled together.

The pooled eluates after FLAG purification were again diluted 4× in KPEM without salt to reduce KCl to ∼ 30 mM and loaded onto a Capto HiRes Q 5/50 GL column (Cytiva). The column was washed overnight with KPEM supplemented with 0.5 mM ATP and 0.5 mM GTP with a flow rate of 0.05 mL/min. Tubulin was eluted with KPEM supplemented with 0.5 mM ATP and 0.5 mM GTP and KCl with concentration ranging between 300 and 400 mM. The eluates were collected and buffer-exchanged into KPEM with a HiPrep 26/20 desalting column (Cytiva). The collected fractions were concentrated in a spin concentrator (Amicon Ultra-15 regenerated cellulose, 30 MWCO) at 3, 234 x g (swing bucket rotor) for about 30 mins.

Tubulin concentrations were determined by absorbance at 280 nm using a Cary 50 spectrophotometer and a molar extinction coefficient of 107,110 M^-1^cm^-1^ for *Homo sapiens* α1bβ4b tubulins and 108 390 M^-1^ cm^-1^ for *Homo sapiens* α1bβ3 tubulins, regardless of wild-type or mutants.

Tubulin was aliquoted in fractions of 20 μL, flash-frozen in liquid nitrogen and stored in the vapour phase of a liquid nitrogen tank.

### Human full-length KIF5B purification

All purification steps were performed at 4°C or on ice, unless otherwise stated. An equal volume of lysis buffer (equilibration buffer supplemented with 10% glycerol, 0.5 mM DTT, 1 mM PMSF, 0.1% CHAPS, ∼250 unit/mL Benzonase® nuclease) was added into the pellets and incubated on ice for about 10 mins. The cells were transferred into a 40 mL douncer and subject to about 40-50 strokes to lyse. This whole cell lysate was clarified by spinning at 200, 000 x g for 35 minutes (Thermo Sceintific, T-865 fixed-angled rotor). A StreptactinXT high capacity FPLC column (IBA Lifesciences, 1 mL) was equilibrated with equilibration buffer (50 mM HEPES, 100 mM potassium acetate, 2 mM MgSO4, 1 mM EGTA, 150 mM NaCl, 100 μM ATP, pH 7.5). The supernatant was loaded onto the column at a flow rate of 0.75 mL/min, then washed with 10 column volume of equilibration buffer at 1 mL/min. KIF5B was eluted with 20 column volume of elution buffer (equilibration buffer + 50 mM biotin) at 0.25 mL/min. Selected fractions were pooled together and subject to buffer exchange into KPEM supplemented with 100 μM of ATP using a PD 10 column (Bio-Rad). The protein was concentrated to desired concentration with a spin concentrator (Amicon Ultra-15 regenerated cellulose, 30 MWCO) at 3, 234 x g (swing bucket rotor) for about an hour. Throughout spinning, the protein was resuspended about every 15 minutes to avoid over-concentration (thus aggregation) at the bottom. Glycerol was added to 20% as cryoprotectant. Aggregates were observed upon addition of glycerol. The aggregates were removed by clarification at 20, 000 x g for 5 mins. Supernatant was aliquoted into 10 μL fractions, flash-frozen in liquid nitrogen and stored in the vapour phase of liquid nitrogen tank.

### Seed preparation

Tubulin was mixed at 1 : 1 : 8 molar ratio fluorescently labelled tubulin : biotinylated tubulin : unlabelled tubulin (porcine brain tubulin, Cytoskeleton) at 26 μM of total tubulin in 1 mM GMPCPP (Jena Bioscience) in buffer KPEM, incubated on ice then divided into 5 μL aliquots and snap-frozen in liquid nitrogen. Seeds were assembled by incubation at 37°C for 30 mins. GMPCPP seeds were pelleted down at 20 psi for 5 minutes at room temperature with an AirFuge (Beckman Coulter). The pellet was gently rinsed with buffer KPEM without nucleotide twice to remove free GMPCPP and gently resuspended in 20 μL KPEM or further diluted to the desired seed concentration for different assays.

### Glass cleaning

Coverslips and glass slides were cleaned by sonication in 3% NeutraCon for 30 minutes, followed by rounds of rinsing and sonication in milliQ water. Slides and coverslips were plasma-cleaned (air plasma, Henniker plasma HPT-200) at 50% power for 5 minutes before use.

### Gliding assays

Tubulin was labelled with 5% Tris-NTA conjugated fluorophores HIS Lite™ OG488-Tris NTA-Ni Complex or HIS Lite™ iFluor® 647 Tris NTA-Ni Complex (AAT Bioquest). Labelling was achieved by mixing tubulin stock with each fluorophore and incubated on ice for 5 mins.

For microtubule assembly in the absence of taxol, iFluor® 647 fluorophore was used for labelling of all tubulin isotypes. 4 μL of tubulin was mixed with 0.5 μL of 10 mM GTP or GDP or KPEM. 2 μL of GMPCPP pig brain seeds was then added into 4 μL of tubulin supplemented with 0.6 μL of 10 mM GTP or GDP or KPEM, the final concentration of tubulin was approximately 20 μM or 40 μM, and then incubated at 37°C for 30 mins in a waterbath.

When taxol was used in microtubule assembly, GMPCPP pig brain MT seeds were omitted. Tris-NTA conjugated Oregon 488 and iFluor® 647 fluorophores were used to label α1bβ3 and α1bβ4b tubulins, wild-type or mutants, respectively. Tubulins at 40 μM were supplemented with 1 mM GTP and incubated at 37°C for 30 mins in a waterbath. An equal volume of 20 μM taxol with 1 mM GTP in KPEM was then added and the mixture incubated for a further 30 mins.

Flow cell channels were built from a larger coverslip and a smaller one with two stripes of double-sided sticky tape as spacers. The surface was passivated with buffer B.2 (0.2 mg/mL α-casein in BRB 80) for 5 minutes. KIF5B was 200× diluted (or 500× diluted when glycerol was not used) in buffer B.2, introduced into the flow cell and incubated for 5 mins. After 5 minutes, unbound kinesin was washed out with gliding buffer (BRB 80 supplemented with 1 mM DTT, 1× GOC, 2 mM ATP and 10 μM taxol or 20% glycerol). Microtubules were suitably diluted (normally 50× when taxol was used in microtubule assembly or 20× in the absence of taxol in which case 20% glycerol was used to stabilise microtubules) in gliding buffer, introduced into the flow cells and incubated for a few minutes to allow sufficient microtubule landing events on kinesin decorated surface. Unbound microtubules were washed out with corresponding gliding buffer.

Imaging of gliding assays was performed at room temperature (between 21-22°C) using a WOSM-TIRF (Warwick Open Source Microscope) with 4-colour light engine, connected to a qCMOS camera (Orca-quest C15550, Hamamatsu) and μManager 2.0. Please see https://wosmic.org/projects/eduwosm/Index.php for details.

Freshly prepared taxol-stabilised α1bβ4b microtubules sometimes got stuck or stuttered on the surface, therefore tracking was done manually with MTrackJ. Only microtubules which showed unimpeded gliding were selected for quantification. The minus-end of each microtubule was tracked every 5 seconds for 35-50 seconds.

### Darkfield/epifluorescence imaging for microtubule dynamics assay

Microtubule dynamics was imaged by darkfield microscope to avoid any impact of fluorescent labelling on the dynamics. Microtubule dynamics were observed by growing microtubules from anchored seeds in a flow cell channel. Flow cell channels were built from two stripes of double-sided tape sandwiched between a 75 mm × 26 mm × 1 mm slide and a 20 mm × 20 mm coverslip, typically with a volume of about 10 μL. Solution exchange was done by flowing in 5 channel volume of reagents. The channel was filled with 0.2 mg/mL PLL-PEG-biotin (SuSoS) in PBS and incubated for 15 minutes, followed by PBS buffer. Subsequently, 1 mg/mL NeutrAvidin (31000, Thermo Scientific) in KPEM was introduced and incubated for 5 minutes, unbound NeutrAvidin was washed out with KPEM. Suitably diluted GMPCPP pig brain seeds were introduced. Unbound seeds were washed out with 1% Tween 20 in KPEM. Tubulins in imaging buffer (1× GOC, 0.5% β-mercaptoethanol and 0.1% tween 20, supplemented with 1 mM GTP and 1 mg/mL BSA in KPEM) were clarified by AirFuge (20 psi for 5 minutes) before being introduced into the channel. The channel ends were sealed with vacuum grease. Imaging was performed at 30°C.

Our darkfield microscope uses an electron-multiplying charge-coupled device (EMCCD) camera (Andor, iXon DU897) fitted to a Nikon E800 microscope with a 100× objective (Plan Fluor NA 0.5-1.3 variable iris, Nikon). The microscope was housed in a custom-built chamber to allow temperature regulation by a heating unit (Air-Therm ATX, World Precision Instruments). Darkfield illumination for unlabelled microtubules was achieved by a combination of a miniature piezo tilt mirror (S-334.1SD, Piezo) reflecting a 532 nm laser (Suwtech Laser) into a circular path and a condenser (Nikon darkfield oil NA 1.43-1.20) to block out unwanted light to generate oblique hollow cone of light. This illuminator was designed and built by Justin Molloy. Epifluorescence illumination for seed visualisation was achieved by a mercury lamp (X-cite exacte, Lumen Dynamics). System control, such as camera exposure time and switching of two illumination sources through shutters, was achieved by μManager 2.0. Images were acquired at 1 frame per sec, taking 1 frame of fluorescence every 600 frames of darkfield images, for 30 minutes. Pixel size was 160 nm.

### Microtubule co-polymerisation assay

Stock tubulins at 40 µM were thawed and supplemented with 1/10 volume 100 μM Tris-NTA conjugated Oregon 488 and iFluor® 647 for α1bβ3 and α1bβ4b tubulins respectively, incubated on ice for 5 minutes. Subsequently, the unbound sites were tagged with 1/10 volume 500 μM of unlabelled Tris-NTA, incubated on ice for 5 minutes. Equimolar amounts of tubulin isotypes were mixed, supplemented with 1/10 volume of 10 mM GTP or just KPEM, and 1/10 volume of GMPCPP pig brain seeds, and incubated at 37°C waterbath for 15 minutes. The microtubules were diluted 10× in 20% glycerol in 1× GOC and 1 mM DTT in BRB80 (without nucleotide) which were introduced into the flow cells decorated with 200× diluted KIF5B as described above for gliding assay.

### Image analysis

Kymographs were generated using ImageJ plugin Kymograph Builder. Stage drift was corrected using “image stabilizer” or “manual drift correction”. The R package “ggplot2” was used to plot the data.

### Primer list

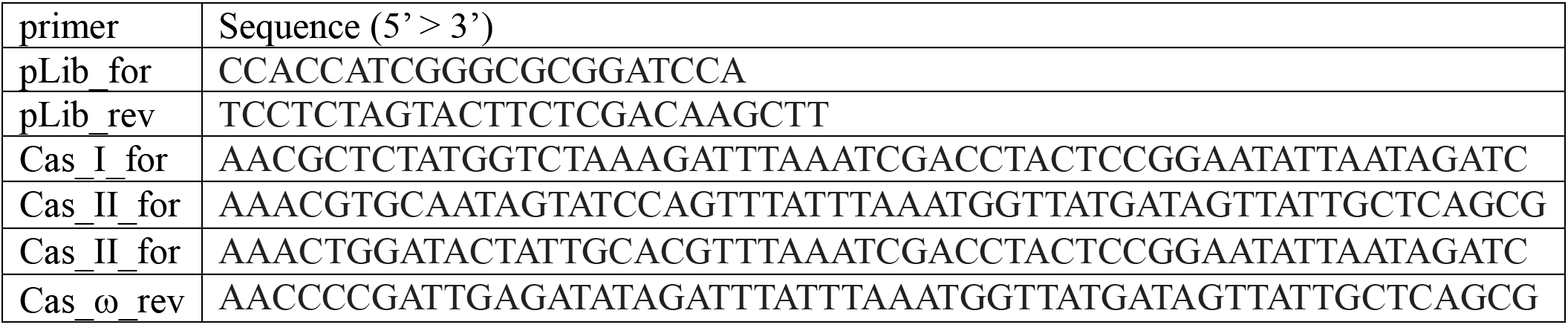

<u>Single underline</u>: linker sequence <u>Double underline</u>: Tag sequence

### β4b tubulin sequences

#### >β4b M-loop binding pocket mutant

MREIVHLQAGQCGNQIGAKFWEVISDEHGIDPSGNYVGDSDLQLERISVYYNEASSH KYVPRAILVDLEPGTMDSVRSGAFGHLFRPDNFIFGQSGAGNNWAKGHYTEGAELV DSVLDVVRKEAESCDCLQGFQLTHSLGGGTGSGMGTLLISKIREEYPDRIMNTFSVVP SPKVSDTVVEPYNATLSVHQLVENTDETYCIDNEALYDICFRTLKLTTPTYGDLNHLV SATMSGVTTCLRFPGQLNADLRKLAVNMVPFPRLHFFMPGFAPLTSRGSQQYRALTV PELTQQMFDAKNMMAACDPRHGRYLTVAAVFRGRMSMKEVDEQMLNVQNKNSSY FVEWIPNNVKTAVCDIPPRGLKMSATFIGNSTAIQELFKRISEQFTAMFRRKAFLHWYT GEGMDEMEFTEAESNMNDLVSEYQQYQDATAEEEGEFEEEAEEEVA<u>GGSGGDYKD DDDK</u>

#### >β4b M-loop mutant

MREIVHLQAGQCGNQIGAKFWEVISDEHGIDPTGTYHGDSDLQLERINVYYNEATGGKYVPRAVLVDLEPGTMDSVRSGPFGQIFRPDNFVFGQSGAGNNWAKGHYTEGAELVDSVLDVVRKEAESCDCLQGFQLTHSLGGGTGSGMGTLLISKIREEYPDRIMNTFSVVPSPKVSDTVVEPYNATLSVHQLVENTDETYCIDNEALYDICFRTLKLATPTYGDLNHLVSATMSGVTTCLRFPGQLNADLRKLAVNMVPFPRLHFFMPGFAPLTSRGSQQYRALTVPELTQQMFDAKNMMAACDPRHGRYLTVAAVFRGRMSMKEVDEQMLNVQNKNSSYFVEWIPNNVKTAVCDIPPRGLKMSATFIGNSTAIQELFKRISEQFTAMFRRKAFLHWYT GEGMDEMEFTEAESNMNDLVSEYQQYQDATAEEEGEFEEEAEEEVA<u>GGSGGDYKD DDDK</u>

#### > β4b WT

MREIVHLQAGQCGNQIGAKFWEVISDEHGIDPTGTYHGDSDLQLERINVYYNEATGGKYVPRAVLVDLEPGTMDSVRSGPFGQIFRPDNFVFGQSGAGNNWAKGHYTEGAELVDSVLDVVRKEAESCDCLQGFQLTHSLGGGTGSGMGTLLISKIREEYPDRIMNTFSVVPSPKVSDTVVEPYNATLSVHQLVENTDETYCIDNEALYDICFRTLKLTTPTYGDLNHLVSATMSGVTTCLRFPGQLNADLRKLAVNMVPFPRLHFFMPGFAPLTSRGSQQYRALTVPELTQQMFDAKNMMAACDPRHGRYLTVAAVFRGRMSMKEVDEQMLNVQNKNSSYFVEWIPNNVKTAVCDIPPRGLKMSATFIGNSTAIQELFKRISEQFTAMFRRKAFLHWYT GEGMDEMEFTEAESNMNDLVSEYQQYQDATAEEEGEFEEEAEEEVA<u>GGSGGDYKD DDDK</u>

### β3 tubulin sequences

#### > β3 M-loop binding pocket mutant

MREIVHIQAGQCGNQIGAKFWEVISDEHGIDPTGTYHGDSDLQLERINVYYNEATGGKYVPRAVLVDLEPGTMDSVRSGPFGQIFRPDNFVFGQSGAGNNWAKGHYTEGAELVDSVLDVVRKECENCDCLQGFQLTHSLGGGTGSGMGTLLISKVREEYPDRIMNTFSVVPSPKVSDTVVEPYNATLSIHQLVENTDETYCIDNEALYDICFRTLKLATPTYGDLNHLVSATMSGVTTSLRFPGQLNADLRKLAVNMVPFPRLHFFMPGFAPLTARGSQQYRALTVPELTQQMFDAKNMMAACDPRHGRYLTVATVFRGRMSMKEVDEQMLAIQSKNSSYFVEWIPNNVKVAVCDIPPRGLKMSSTFIGNSTAIQELFKRISEQFTAMFRRKAFLHWYT GEGMDEMEFTEAESNMNDLVSEYQQYQDATAEEEGEMYEDDEEESEAQGPK<u>GGSG GDYKDDDDK</u>

#### > β3 M-loop mutant

MREIVHIQAGQCGNQIGAKFWEVISDEHGIDPSGNYVGDSDLQLERISVYYNEASSHKYVPRAILVDLEPGTMDSVRSGAFGHLFRPDNFIFGQSGAGNNWAKGHYTEGAELVDSVLDVVRKECENCDCLQGFQLTHSLGGGTGSGMGTLLISKVREEYPDRIMNTFSVVPSPKVSDTVVEPYNATLSIHQLVENTDETYCIDNEALYDICFRTLKLTTPTYGDLNHLVSATMSGVTTSLRFPGQLNADLRKLAVNMVPFPRLHFFMPGFAPLTSRGSQQYRALTVPELTQQMFDAKNMMAACDPRHGRYLTVATVFRGRMSMKEVDEQMLAIQSKNSSYF VEWIPNNVKVAVCDIPPRGLKMSSTFIGNSTAIQELFKRISEQFTAMFRRKAFLHWYT GEGMDEMEFTEAESNMNDLVSEYQQYQDATAEEEGEMYEDDEEESEAQGPK<u>GGSG GDYKDDDDK</u>

#### > β3 WT

MREIVHIQAGQCGNQIGAKFWEVISDEHGIDPSGNYVGDSDLQLERISVYYNEASSHKYVPRAILVDLEPGTMDSVRSGAFGHLFRPDNFIFGQSGAGNNWAKGHYTEGAELVDSVLDVVRKECENCDCLQGFQLTHSLGGGTGSGMGTLLISKVREEYPDRIMNTFSVVPSPKVSDTVVEPYNATLSIHQLVENTDETYCIDNEALYDICFRTLKLATPTYGDLNHLVSATMSGVTTSLRFPGQLNADLRKLAVNMVPFPRLHFFMPGFAPLTARGSQQYRALTVPELTQQMFDAKNMMAACDPRHGRYLTVATVFRGRMSMKEVDEQMLAIQSKNSSYFVEWIPNNVKVAVCDIPPRGLKMSSTFIGNSTAIQELFKRISEQFTAMFRRKAFLHWYT GEGMDEMEFTEAESNMNDLVSEYQQYQDATAEEEGEMYEDDEEESEAQGPK<u>GGSG GDYKDDDDK</u>

### α1b tubulin

#### > α1b WT

MRECISIHVGQAGVQIGNACWELYCLEHGIQPDGQMPSDKTIGGGDDSFNTFFSETGAGKHVPRAVFVDLEPTVIDEVRTGTYRQLFHPEQLITGKEDAANNYARGHYTIGKEIIDLVLDRIRKLADQCTGLQGFLVFHSFGGGTGSGFTSLLMERLSVDYGKKSKLEFSIYPAPQVSTAVVEPYNSILTTHTTLEHSDCAFMVDNEAIYDICRRNLDIERPTYTNLNRLISQIVSSITASLRFDGALNVDLTEFQTNLVPYPRIHFPLATYAPVISAEKAYHEQLSVAEITNACFEPANQMVKCDPRHGKYMACCLLYRGDVVPKDVNAAIATIKTKRSIQFVDWCPTGFKVGINYQPPTVVPGGDLAKVQRAVCMLSNTTAIAEAWARLDHKFDLMYAKRAF VHWYVGEGMEEGEFSEAREDMAALEKDYEEVGVDSVEGEGEEEGEEY<u>GGSGGHH HHHHHH</u>

